# Yeast EndoG prevents genome instability by degrading cytoplasmic DNA

**DOI:** 10.1101/2023.12.13.571550

**Authors:** Yang Yu, Xin Wang, Jordan Fox, Ruofan Yu, Pilendra Thakre, Brenna McCauley, Nicolas Nikoloutsos, Qian Li, P. J. Hastings, Weiwei Dang, Kaifu Chen, Grzegorz Ira

**Author notes:** equal contribution (Yang Yu, Xin Wang, Jordan Fox). Correspondence: Grzegorz Ira, Kaifu Chen.

## Abstract

In metazoans release of mitochondrial DNA or retrotransposon cDNA to cytoplasm can cause sterile inflammation and disease. Cytoplasmic nucleases degrade these DNA species to limit inflammation. It remains unknown whether degradation these DNA also prevents nuclear genome instability. To address this question, we decided to identify the nuclease regulating transfer of these cytoplasmic DNA species to the nucleus. We used an amplicon sequencing-based method in yeast enabling analysis of millions of DSB repair products. <u>Nu</u>clear <u>mt</u>DNA (NUMTs) and retrotransposon cDNA insertions increase dramatically in nondividing stationary phase cells. Yeast EndoG (Nuc1) nuclease limits insertions of cDNA and transfer of very long mtDNA (>10 kb) that forms unstable circles or rarely insert in the genome, but it promotes formation of short NUMTs (∼45-200 bp). Nuc1 also regulates transfer of cytoplasmic DNA to nucleus in aging or during meiosis. We propose that Nuc1 preserves genome stability by degrading retrotransposon cDNA and long mtDNA, while short NUMTs can originate from incompletely degraded mtDNA. This work suggests that nucleases eliminating cytoplasmic DNA play a role in preserving genome stability.

---

Mitochondria and chloroplasts carry multiple copies of a small genome, and transfer of DNA from these organelles to the nuclear genome is a ubiquitous and ongoing process. Genomes of most eukaryotes contain many fragments from these organelles ^1, 2^. Insertions of these DNA species at nuclear genome can inactivate genes, form new exons, act as new or modify existing origins of replication, modify gene expression, and impact adaptation and genome evolution ^3–7^. Insertions of mtDNA and cDNA form by their capture at spontaneous, programmed or induced DNA double-strand breaks (DSB) via nonhomologous end joining (NHEJ) (e.g.^8–12^). The frequency of these insertion events increases with stress, aging, and tumorigenesis ^8, 13–17^, yet little is known about the enzymes that regulate their transfer to the nucleus. While it is established that nucleases such as EndoG and TREX1 can degrade cytoplasmic DNA in humans to prevent activation of the immune response^18, 19^, it remains unclear whether these mechanisms could also play a role in preventing cytoplasmic DNA from causing harmful genome instability. A major barrier to addressing this question is the difficulty of detecting these rare insertion events. Here we introduce a high-throughput amplicon sequencing pipeline that enables detection and analysis of many thousands of these events in yeast and we provide evidence that the yeast EndoG homolog prevents genome instability in stressed cells by degrading cytoplasmic DNA species.

## *Break-Ins* method for insertion analysis

To study nuclear genome instability caused by insertion of mtDNA and retrotransposon cDNA, we developed a high-throughput amplicon sequencing based method called *Break-Ins* (*<u>Break Ins</u>ertions*) which allows screening of hundreds of thousands of NHEJ products simultaneously for the events carrying NUMTs (**Fig. 1a**). This method is suitable for testing many mutant strains or conditions simultaneously. We used yeast haploid cells carrying a galactose-inducible HO endonuclease that generates a single double-strand break (DSB) per genome at the *MAT***a** locus. In this strain homologous sequences for DSB repair, *HML* and *HMR,* were deleted. Therefore, this DSB can only be repaired by NHEJ; only the cells that repaired the break imprecisely, altering the HO recognition site, can survive continuous HO induction and form colonies ^20^. To identify rare long >10 bp insertion events, repaired *MAT***a** loci were amplified by PCR using primers located upstream and downstream from the HO cut site. Libraries of PCR fragments were prepared using a modified Illumina procedure and subjected to amplicon sequencing using the MiSeq platform (**Fig. 1a-b**; **Supplementary Fig. 1a-e**). To test this method, we initially compared wild-type cells in which insertions are very rare and Dna2-deficient cells in which insertions are common ^21^. Over 100,000 colonies corresponding to independent NHEJ repair events were pooled in wild-type or Dna2-deficient cells. Amplification of the *MAT***a** locus with NHEJ products from Dna2-deficient cells, but not from WT cells, showed a smear of products above the band corresponding to the normally repaired *MAT***a** fragment size, suggesting many insertions in mutant but not WT cells (**Fig. 1c**). Only reads carrying insertions of >10 bp, hereafter called “insertions”, were subjected to analysis. In wild-type growing cells, only 8 unique insertions were identified, three of which were derived from Ty retrotransposons, two from rDNA, and three from other parts of the nuclear genome (**Supplementary Table 1**). Overall, a ∼280-fold increase in insertions was observed in *dna2*Δ *pif1-m2* cells (**Supplementary Fig. 2a**). Repeating amplicon sequencing of the same DNA but using primers located further from the DSB led to identification of ∼25% new insertions (**Supplementary Fig. 2b**). Nearly all unique insertions observed only with one pair of primers were represented by a lower read number, likely representing events more difficult to sequence. The distribution of inserted DNA within the genome, features of DNA inserted, and insertion junctions were comparable to previous Sanger analysis of individual NHEJ products (**Supplementary Fig. 2c-e**) ^21^. Together we conclude that *Break-Ins* produces high-quality data and can be applied to study the transfer of cytoplasmic DNA to nucleus.

**Figure 1.**
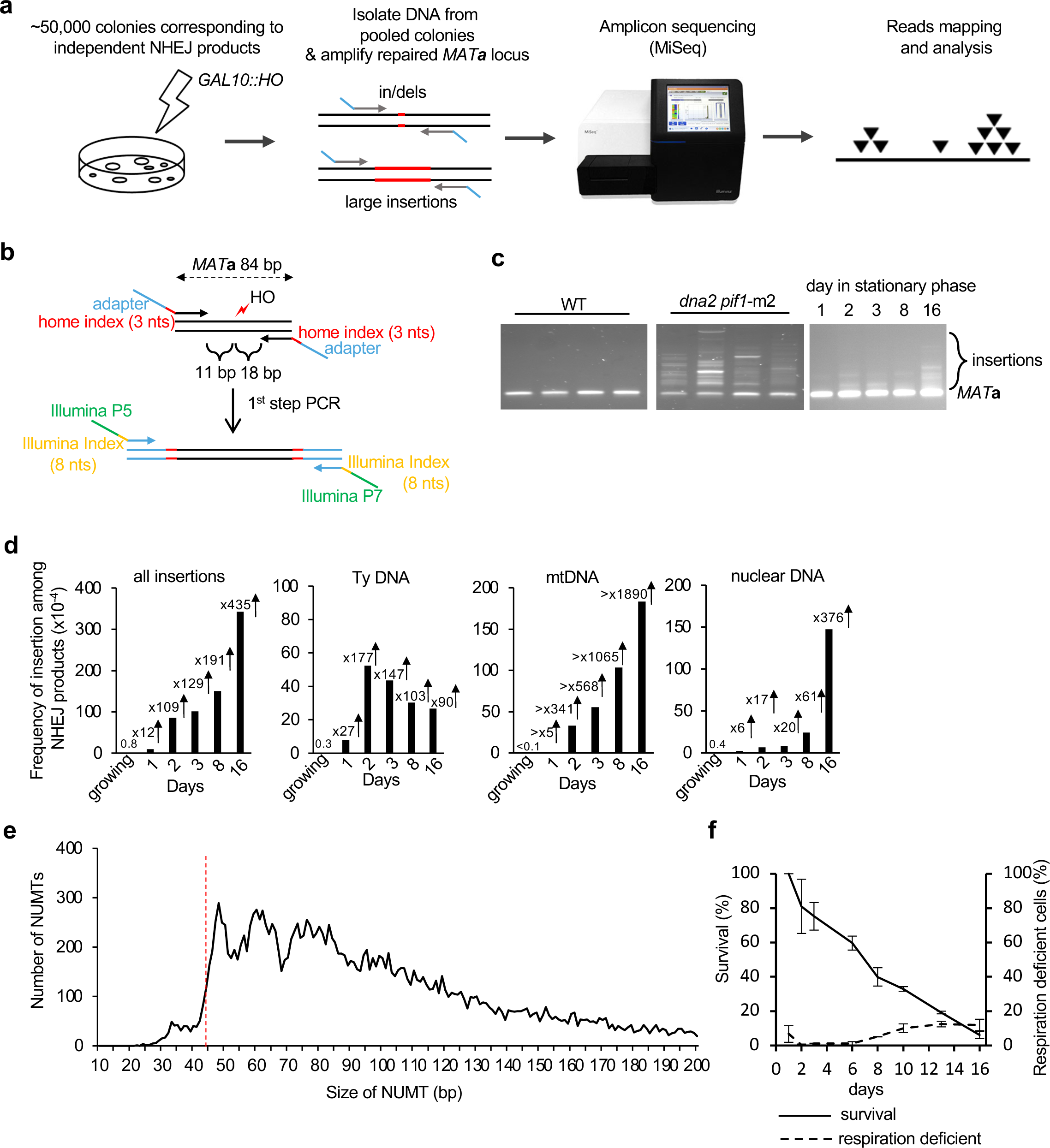
Insertions of cytoplasmic DNA species in stationary phase cells. **a,** Schematic of the *Break-Ins* method. **b**, Schematic of PCR amplification of the repaired *MAT* locus with two sets of indexes. Two sets of indexes were used to limit incorrect sample assignment by index hopping. **c**, PCR amplification of repaired *MAT* locus in WT, *dna2*Δ *pif1-*m2 mutant and stationary phase cells. **d**, Frequency and types of DNA inserted at DSBs in wild-type stationary phase cells. “all insertions” include single and complex events. Minimal increase indicated for mtDNA as there was no single mtDNA in growing cells. **e**, Insertion size analysis of NUMTs. The insertions originating from mtDNA were from 16 days stationary phase cells (n=21,348). Red dashed line indicates minimal size of DNA for efficient Ku70/80 heterodimer binding ^26^. **f**, Survival and respiration proficiency of cells plated on rich media after 1-16 days in stationary phase, (mean ± SD; n=3).

## Frequent NUMTs and retrotransposon cDNA insertions in stationary phase cells

Analysis of insertions in wild-type cells suggests that the transfer of mtDNA or cDNA is rare in dividing cells grown in optimal conditions (**Fig. 1d**). To study the transfer of cytoplasmic DNA to the nucleus, we decided to test NHEJ products from nondividing, stationary phase cells where transfer of mtDNA was observed in fission yeast ^8^ and where retrotransposon transcription increases dramatically ^22^. NHEJ products were analyzed in nondividing cells incubated in synthetic complete media for 1, 2, 3, 8 or 16 days followed by DSB induction on YEPGal plates. The time of survival after cells enter stationary phase due to the exhaustion of nutrients is often referred to as chronological lifespan ^23^. PCR amplification of the *MAT***a** locus suggests a high level of inserted DNA (**Fig. 1c**). Indeed, a dramatic increase in insertions was observed in early 1-, 2- or 3-days stationary phase wild-type cells (**Fig. 1d**). At these earliest time points (1-3 days), inserted DNA originated mostly from mtDNA (∼340-570-fold increase at 2-3 days) and Ty retrotransposons (∼150-fold increase at 2-3 days). At later 8- and 16-day timepoints, NUMTs further increased up to over 1900-fold compared to growing cells and pieces of the nuclear genome were inserted as well (∼60-380-fold increase). Insertions from all DNA sources (Ty, mtDNA, nuclear genome) were dependent on Lig4, essential for NHEJ, and largely dependent on DNA polymerase Pol4 known to promote error-prone NHEJ ^24^. They were independent of Rad51 or a nonessential component of Polδ, Pol32, which makes it unlikely that break-induced replication-like events were involved (**Supplementary Fig. 3a-b**). Very rare insertions were observed in *pol4*Δ cells (**Supplementary Table 1**). Genetic requirements and microhomology at DSB ends (**Supplementary Fig. 3c**) suggest that most of the insertions occurred by capture of DNA fragments at DSB by NHEJ. Additional analysis of ∼0.5 mln NHEJ products from 16-day stationary phase wild-type cells revealed together over 20,000 NUMTs. The size of NUMTs ranged from 22 to 492 bp, comparable to natural yeast NUMTs (**Supplementary Fig. 3d**, ^25^). Interestingly, the minimal size of mtDNA for efficient insertion is 44-46 bp (**Fig 1e**) which likely reflects the DNA length-dependent cooperative binding of Ku70/Ku80 heterodimer to DNA, where ∼45 bp-long dsDNA is the minimal DNA size for optimal binding ^26^. The size of insertions originating from Ty retrotransposons was comparable to NUMTs while the size of insertions from the nuclear genome was about twice as long (**Supplementary Fig. 3d)**. Among NHEJ products that have altered the HO cleavage site (∼0.1-0.3% of cells plated), about 1% carried NUMTs which corresponds to about one NUMT per 10^5^ cells plated. Inserted fragments of the nuclear genome often originated from fragile regions of the genome such as telomeric, R-loop-prone, or repetitive regions (**Supplementary Fig. 3e**). Thus, insertions at DSBs can occur in stressed stationary phase wild-type cells, and are not limited to mutant cells such as Dna2-deficient yeast or human cells, or cancer cells ^21, 27, 28^.

Insertions of mtDNA at DSBs could originate from other cells in culture, dead or alive, or from loss of respiration coupled with loss of mtDNA. The level of respiration deficient petite colonies that either lost (*rho*^0^) or severely mutated mtDNA (*rho*^-^) does not change in day 1-3 of stationary phase as measured by ability to grow on a nonfermentable carbon source (**Fig. 1f**). Also, viability in days 1-3 is only slightly decreased while the number of NUMTs increases dramatically (**Fig. 1f**). Thus, NUMT formation in early stationary phase cells is not associated with overall loss of mtDNA or loss of viability. Another argument against dead cells being a source of mtDNA for NUMT formation is that *sch9Δ* mutant maintains 100% viability at 8 days stationary phase (**Supplementary Fig. 3f**) ^29^, yet the number of insertions is similar when compared to WT. Finally, mtDNA fragments could originate from a cell’s own mitochondria, including connected mother-daughter pairs, or from other live cells in culture. We favor the first possibility because a previous study demonstrated that mtDNA can transfer to the nucleus but not between two yeast strains ^30^. Overall, we conclude that NUMT formation increases in stationary phase cells and that mtDNA inserted at DSBs likely originates from within the same cell.

## Nuc1 nuclease regulates cDNA and mtDNA insertions

In humans nucleases such as EndoG eliminate cytoplasmic DNA to prevent inflammation (review in ^31^) but it remains unknown if they impact nuclear genome stability. To test this in yeasts we followed genomic insertions of mtDNA and cDNA in cells lacking Nuc1, homolog of human EndoG. The primary localization of Nuc1 is in mitochondria but during stress and sporulation it can relocate to the cytoplasm ^32, 33^. EndoG nucleases have DNA endonuclease and 5’ exonuclease activities with preference for ssDNA over dsDNA ^34^. *NUC1* deletion does not affect DNA insertions at DSBs in growing cells (**Supplementary Table 1**); however, we found it has a profound impact in stationary phase cells. In *nuc1*Δ cells, the overall level of insertions is higher due to ∼Δ-fold increase of retrotransposon cDNA insertions. This is consistent with the possibility that Nuc1 degrades Ty1 cDNA that forms in cytoplasm. Unexpectedly, NUMTs were reduced by ∼12-folds in *nuc1*Δ suggesting a more complex role of this nuclease in regulation of mtDNA transfer to nucleus that is addressed below. Insertions from the nuclear genome at 8 days remained at the level comparable to wild type (**Fig. 2a**). Nuc1 nuclease activity is responsible for the changes of NUMTs and Ty cDNA insertions, as evidenced by a similar pattern of insertions in *nuc1*-H138A nuclease dead mutant and *nuc1*Δ cells (**Fig. 2a**). Considering the broad impact of Nuc1 on generation of insertions, we analyzed the pattern of insertions additionally at 3 and 16 days and observed a comparable effect on mtDNA and Ty DNA (**Fig. 2b**). Low mtDNA insertions in *nuc1*Δ cells are not related to a change in viability or respiration proficiency (**Fig. 2c**). By 16 days, NUMTs increased in *nuc1*Δ cells, indicating the existence of a less efficient Nuc1-independent pathway of NUMT formation of slightly longer size.

**Figure 2.**
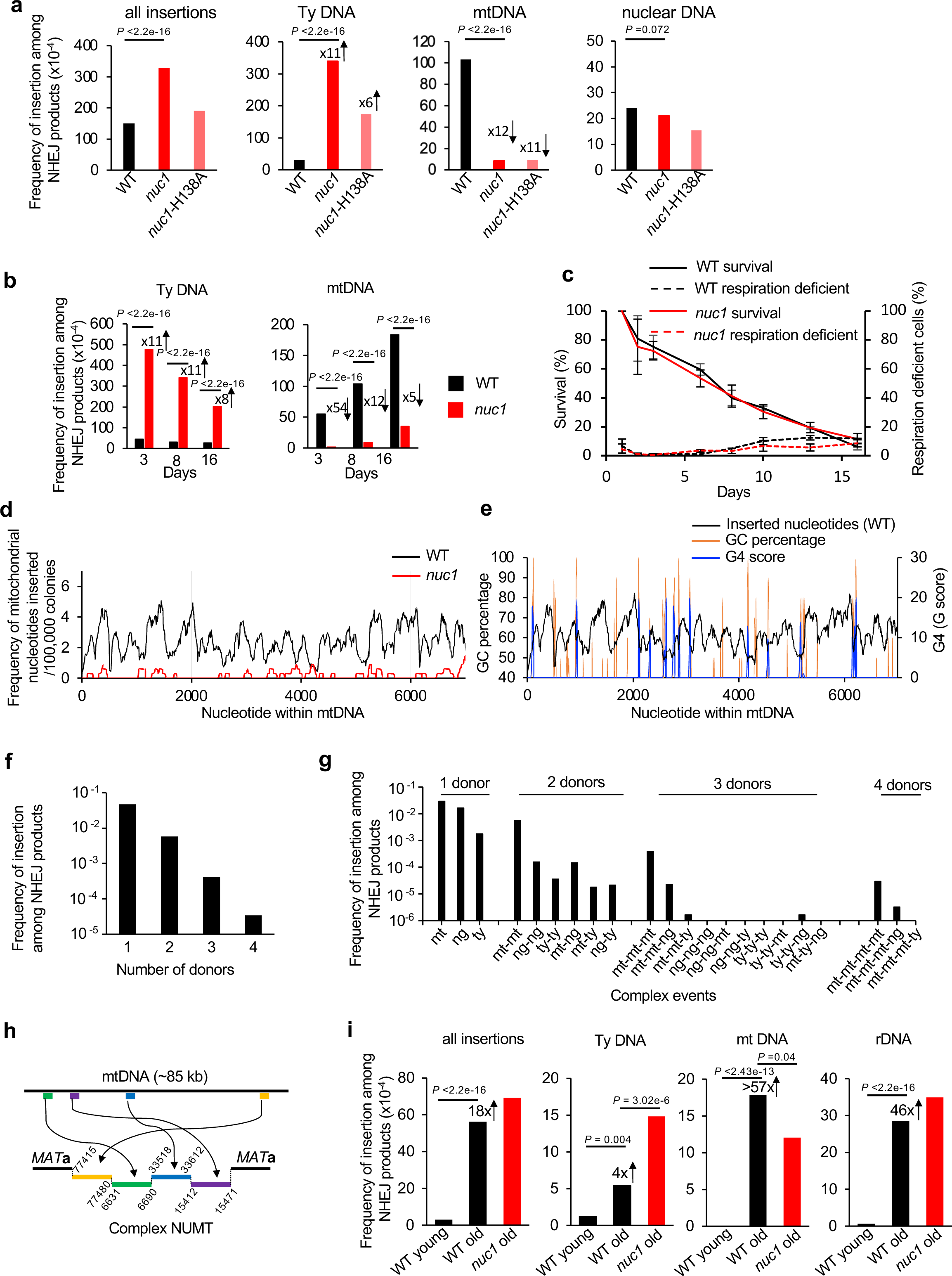
Nuc1 regulates genome instability caused by cytoplasmic DNA. **a-b**, Frequency and types of DNA inserted at DSBs in wild-type, *nuc1*Δ, and *nuc1*-H138A mutant cells in 8 days (a) or 3, 8 and 16 days (b) stationary phase cells. P values were determined using χ^2^ test. n – number of NHEJ products tested shown in Supplementary Table 1. **c**, Survival and respiration proficiency of wild-type and *nuc1*Δ cells plated on rich media after 1-16 days in stationary phase, (mean ± SD; n= 3). **d**, Analysis of sequences inserted at DSB from the first 7 kb of mtDNA in wild-type and *nuc1*Δ cells (complete mtDNA sequence shown in Supplementary Figure 4). Single donor insertions were analyzed. **e**, Sequences inserted into DSBs from the first 7 kb of mtDNA and the distribution of G4 forming sequences and GC content within the first 7 kb of mtDNA. Single donor insertions were analyzed. **f**, Analysis of complex insertions in wild-type cells at 16 days. Number of insertions involving 1, 2, 3 or 4 DNA pieces are shown. **g**, Distribution of different complex insertion events in wild-type at 16 days. Mt-mitochondrial DNA, ty-retrotransposon DNA, ng-nuclear genome DNA. **h**, Example of a complex NUMT. **i**, Frequency and types of insertions in wild-type and *nuc1*Δ cells in young cells and aged mother cells. P values were determined using χ^2^ test. n - number of NHEJ products tested shown in Supplementary Table 1.

NUMTs in wild-type cells originated from throughout the 86 kb-long yeast mitochondrial chromosome (**Fig. 2d, Supplementary Fig. 4a**). We have not observed insertions of spliced mtDNA fragments, indicating that reverse transcribed RNA is not a source of inserted mtDNA. Interestingly, G-rich sequences often mark the boundaries of the peaks of inserted mtDNA (**Fig. 2e**). Indeed, insertions from wild-type cells (p-value < 2.2e-16, one-sided Wilcoxon test) but not *nuc1*Δ (p-value = 1, one-sided Wilcoxon test) have an inverse relationship with G4- or GC-rich sequences. Nuc1 homolog EndoG preferentially binds and cuts poly-G rich sequences including G4 structure forming sequences ^35–37^. Additionally, yeast Nuc1 and human EndoG are responsible for mtDNA deletions, which often occur within or at the boundaries of G sequence clusters (e.g.^35^). Thus, possible preferential mtDNA degradation of G rich sequences by EndoG nucleases could reflect a slight insertion bias toward AT rich sequences. However, the nuclease activity of yeast Nuc1 toward G4 structures has not yet been studied.

Some of the sequenced NUMTs within eukaryotic genomes are complex, with multiple fragments joined together or in combination with pieces of nuclear genome or Ty retrotransposons (e.g.^13, 38, 39^). Among ∼33,000 insertions obtained from 16-day stationary phase wild-type cells, nearly 4,000 contain complex insertions of two to four different fragments joined together (**Fig. 2f**). The frequency of complex NUMTs, but not of other multi-insertions, is higher than expected by chance (p-value of 0.0002, Fisher’s exact test), a phenomenon similar to the higher than expected level of multiple *de novo* NUMTs observed in cancer cells ^13^. A possible explanation for the high level of multi-insertions is unequal release of mtDNA among cells. Most of these complex events carried multiple fragments of mtDNA, while the rest included only Ty DNA or nuclear DNA or were the mixture of multiple DNA sources (**Fig. 2g**). An example of a complex NUMT is shown in **Fig. 2h**. Complex mitochondrial rearrangements are reminiscent of chromothripsis events, in which fragments of nuclear chromosomes are subjected to clustered rearrangements ^40^. As suggested previously, shattering of the mitochondrial genome followed by random assembly of DNA fragments could constitute a mito-chromothripsis ^13^. Our data provide compelling evidence for such a possibility with Nuc1 being responsible for mtDNA fragmentation.

NUMTs increase during aging across many species ^14, 41, 42^. To test the transfer of cytoplasmic DNA to nucleus during aging and a possible role of Nuc1, we first analyzed the lifespan of the strains used to capture DNA sequences at DSB using a microfluidic device. Wild-type and *nuc1Δ* showed comparable lifespans (**Supplementary Fig. 4e**). We then isolated old mother cells corresponding to ∼60-80% of their lifespan. Old mother cells were plated on YEPGal plates to induce a DSB and repair products were followed using *Break-Ins* analysis as described above. An 18-fold increase of insertions was observed compared to young cells. Inserted DNA originated from mtDNA (>57-fold increase), Ty retrotransposon (4-fold increase) and the genome, mostly from rDNA (**Fig. 2i**). Further, we tested the possible impact of Nuc1 on NUMTs and Ty insertions in aged cells. In *nuc1*Δ cells we observed a small (less than 2-fold) reduction of NUMTs and 3-fold increase of Ty cDNA insertions (**Fig. 2i**). A total of 158 NUMTs were observed in aged WT or *nuc1*Δ cells and they originated from throughout the mtDNA genome (**Supplementary Fig. 4b**). Thus, Nuc1 has a similar role during aging but additional nuclease(s) can fragment mtDNA.

## Nuc1 degrades long mtDNA to prevent its transfer to nucleus

*Break-Ins* method can only monitor transfer/insertions of short mtDNA. To test a possible role of Nuc1 in transfer of long mtDNA we used a previously established assay ^43^. In this assay, transfer of long mtDNA fragments carrying a *TRP1* reporter gene that is not expressed in mitochondria to the nucleus generates cells expressing *TRP1* and Trp^+^ colonies. Nuclear mtDNAs are maintained as unstable circular fragments of few to >30 kb that are easily lost and thus do not represent stable NUMTs ^43^. The multitude of putative origins of replication within the AT-rich mitochondrial genome allows propagation of mtDNA as circular DNA ^5, 7^. Unexpectedly, and in contrast to short NUMTs that are nearly eliminated in *nuc1*Δ mutant cells, transfer of large mtDNA was greatly increased upon loss of *NUC1* (**Fig. 3a, b**). A 4- to 22-fold increase of mtDNA transfer frequency was observed in *nuc1*Δ stationary phase cells when compared to wild-type cells. We confirmed that nuclear *TRP1*-mtDNA forms circles by testing these DNA by Southern blot in cells from which mtDNA was eliminated by EtBr treatment as described previously (**Supplementary Fig. 5**)^43^. As expected, a comparable increase was noted in *nuc1*-H138A nuclease dead cells (**Fig. 3a, b**). At 8 days ∼1% of *nuc1*Δ cells carried nuclear *TRP1*-mtDNA, which is likely an underestimate of total mtDNA transfer, considering that we can only follow *TRP1*-marked mtDNA fragments. It is ∼1000-fold higher level of mtDNA transfer to the nucleus when compared to formation of short stable NUMTs at HO breaks in wild-type cells. Thus, the release of mtDNA from mitochondria is independent of Nuc1 status. As Nuc1 is known to relocate to the cytoplasm in stressed cells ^32, 33^, we propose that cytoplasmic Nuc1 prevents frequent long mtDNA transfer to the nucleus by degrading mtDNA released from the mitochondrion, while incomplete degradation of long mtDNA by Nuc1 may result in shorter mtDNA fragments that form rare NUMTs (model in **Fig. 3c**). These results also suggest that having long mtDNA transferred to the nucleus is not a common prerequisite of short stable NUMTs, as the number of NUMTs is greatly decreased in *nuc1*Δ, yet long mtDNA transfers more frequently.

**Figure 3.**
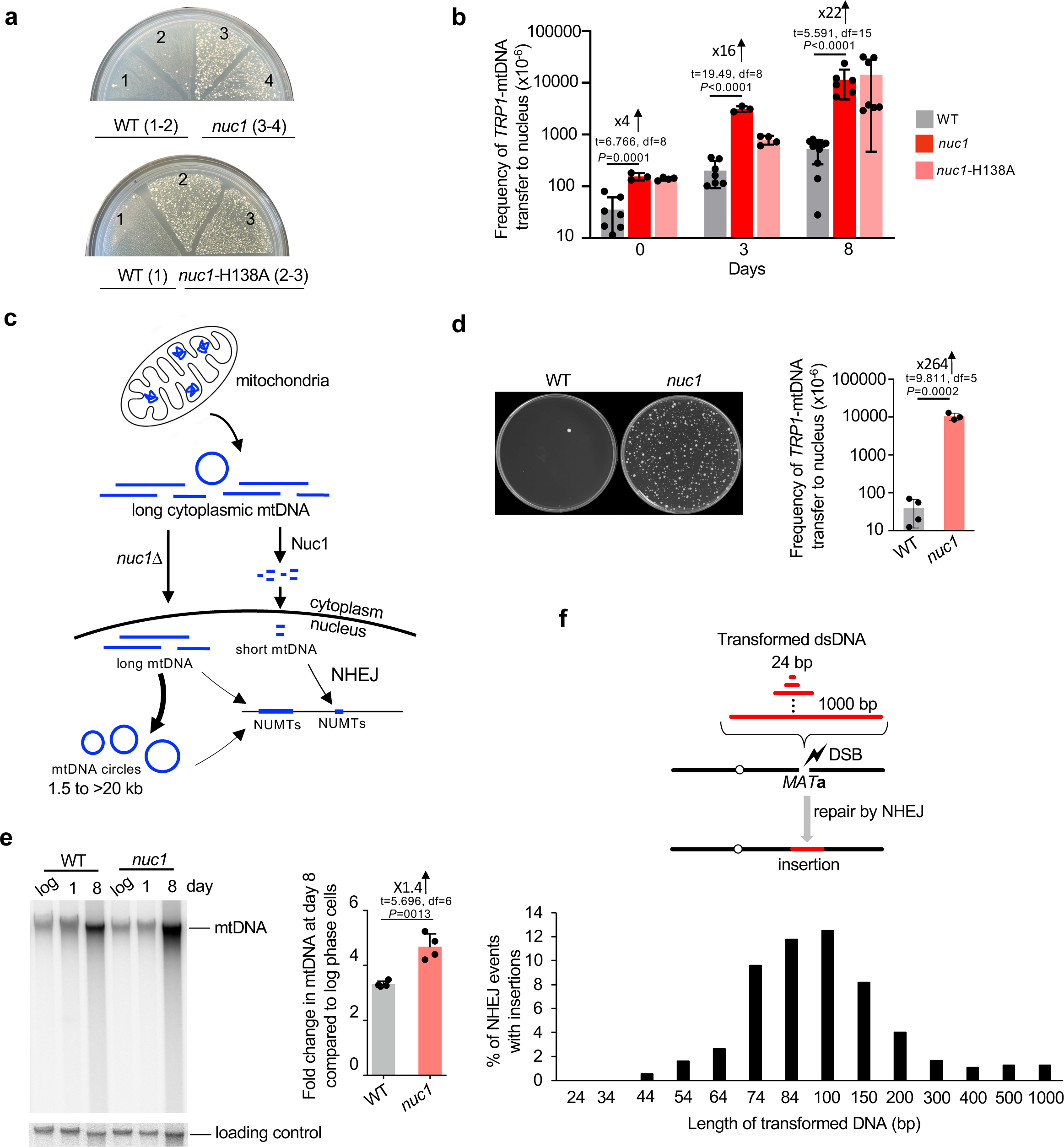
Nuc1 suppresses transfer of long mtDNA to nucleus. **a**, Analysis of *TRP1*-mtDNA transfer to nucleus. Wild-type, *nuc1*Δ and *nuc1*-H138A mutants cells replica plated from rich YEPD media onto plates with media lacking tryptophan. Trp^+^ colonies represent transfer of *TRP1*-mtDNA to nucleus. **b**, Frequency of Trp^+^ colonies in wild-type and *nuc1*Δ cells, (mean ± SD; n≥3; P values determined using unpaired two-tailed t-test). **c**, Model of Nuc1 mediated control of mtDNA transfer to the nucleus. In stationary phase cells, mtDNA is released from mitochondria in Nuc1-independent manner. It is degraded in cytoplasm by Nuc1 nuclease. Incomplete degradation leaves small mtDNA pieces that can be easily integrated at DSBs. In the absence of Nuc1, mtDNA is not degraded and large mtDNA pieces are transferred to the nucleus where they either form circular mtDNA of different sizes or rarely integrate in the genome. **d**, Random spore analysis on plates with media lacking tryptophan. Trp^+^ colonies represent transfer of *TRP1*-mtDNA to nucleus. Frequency of spores carrying nuclear *TRP1*-mtDNA is shown. (mean ± SD; n≥3; P values determined using unpaired two-tailed t-test). **e**, Southern blot analysis and amount of mtDNA in wild-type and *nuc1*Δ at 8 days stationary phase cells, (mean ± SD; n = 4; P values determined using unpaired two-tailed t-test). **f**, Scheme and analysis of insertion originating from transformed DNA fragments of different sizes. Frequency of insertions of annealed oligonucleotides (24-74bp) or PCR fragments (84-1000bp) are shown. 480 to 1680 independent NHEJ products were tested by PCR.

Next, we decided to test whether Nuc1 function in limiting mtDNA transfer to nucleus also applies to yeast stationary phase diploid cells that undergo meiotic recombination and sporulation. Wild-type and *nuc1*Δ*/nuc1*Δ diploids were sporulated and subjected to random spore analysis on plates lacking tryptophan. As shown in **Fig. 3d**, spores produced in Nuc1-deficient cells showed a dramatic >250-fold increase of Trp^+^ colonies corresponding to mtDNA transfer to the nucleus. About 1% of all spores carried nuclear *TRP1* marked mtDNA in Nuc1-deficient cells. Thus, Nuc1 controls the frequency of mtDNA transfer to nuclei in diploid cells during meiosis. 180 independent Trp^+^ spores were tested for the maintenance of nuclear *TRP1*-mtDNA and only one carried a stable *TRP1* marker, indicating that long nuclear mtDNA fragments also integrate into the genome. In summary, Nuc1 appears to prevent the transfer of long mtDNA to the nucleus during sporulation that can lead to rare stable genomic integration.

As Nuc1 appears to control the size and frequency of mtDNA transfer to the nucleus, we compared the level of mtDNA in wild-type and *nuc1*Δ cells cultured in stationary phase for 8 days. Isolated DNA was separated on agarose gel and subjected to Southern blot analysis using a mtDNA probe. Despite decreasing nutrients, overall mtDNA copy number per cell increased by 3-4-fold in 8-day-old stationary phase cells. Such increase was previously observed and could be related to a manyfold increase of mitochondria volume in stationary phase wild-type cells ^44, 45^. There is a 1.4-fold increase of mtDNA in *nuc1*Δ cells when compared to wild type, likely reflecting mtDNA that Nuc1 would degrade, preventing its transfer to the nucleus (**Fig. 3e**).

In *nuc1*Δ cells, long mtDNA is transferred to the nucleus, yet this DNA is not frequently inserted at HO breaks or anywhere else in the genome. This means that the short size of NUMTs is not likely related to the inability of longer DNA to cross the nuclear membrane. Also, MiSeq read length of up to 600 bp cannot explain a clear bias toward ∼100 bp NUMTs and far fewer NUMTs of 200-600 bp. One possibility is that NHEJ has a preference for inserting shorter DNA. To test this, we transformed an equal molar amount of 14 DNA fragments ranging from 24 bp to 1 kb, each carrying the same sequences on 5’ and 3’ ends matching HO cut overhangs. Each fragment was transformed independently, and cells were spread on YEPGal to induce DSB. To avoid any Illumina sequencing bias toward DNA fragments of a particular size, we tested 400 to >1000 NHEJ products individually by PCR. The shortest fragment inserted was 44 bp long and fragments of ∼60-70 bp to ∼150 bp inserted most frequently (**Fig. 3f**). The length of transformed DNA optimal for insertion at DSB is comparable to the size of all types of inserted DNA (mtDNA, Ty DNA, nuclear genome); however, NUMTs and Ty DNA insertions are on average slightly shorter while insertions from the nuclear genome are longer when compared to the size of insertions of transformed DNA (**Supplementary Fig. 3d**). Thus, the bias toward short insertions (∼45 to ∼300 bp) could stem from easier insertion of short DNA fragments at DSBs by NHEJ. In general, the size of insertions observed in different experimental systems and species is within the range observed here ^9, 12, 21, 39^.

## Nuc1 degrades retrotransposon cDNA

In stationary phase cells, insertions of Ty DNA increase by ∼90-180-fold and Nuc1 absence leads to a further increase by ∼Δ-fold (**Fig. 1d, 2a**). To analyze transposon insertions in detail we selected the ones originating from Ty1, the most common retrotransposon in yeast. All regions of Ty1 contributed to insertions, and as previously shown the 3’ LTR region is a clear hotspot in wild-type cells (**Fig. 4a, Supplementary Fig. 4c**) ^10, 11, 46^. In *nuc1*Δ cells, a large increase of inserted Ty1 fragments was observed at the region outside of LTR (∼10 to 40-fold), particularly from the primer binding site (PBS) to the central priming site (PPT2), while the increase of the 3’ LTR region was very modest (∼2-fold). A large fraction of 3’ LTR insertions in wild-type cells (40/56) carried extra nucleotides present only in cDNA, consistent with intermediates of Ty1 replication being inserted at DSBs ^11, 46^. Increased insertions of Ty DNA in stationary phase cells are consistent with greatly increased Ty transcription ^22^. However, high Ty1 expression is not associated with change in transposition in wild-type or *nuc1*Δ old stationary phase cells as measured using a previously described Ty1 transposition reporter assay ^47^ (**Supplementary Fig. 4d**). The full length Ty cDNA amount was measured by Southern blot and shows no major difference between wild-type and *nuc1*Δ cells that could account for the dramatic increase of cDNA insertions observed in *nuc1*Δ (**Fig. 4b**). As expected, no cDNA was observed in the absence of Spt3 needed for Ty1 expression (**Fig. 4b**). Besides complete linear or circular cDNA of Ty1, another possible source of fragments inserted at DSBs could be intermediates of Ty replication. ssDNA minus strand intermediates of Ty1 replication can be inserted at DSBs and can recombine with the yeast genome ^48^, and nucleocapsids are known to carry Ty1 replication intermediates observed as a smear of Ty1 cDNA ^49^. We found ∼7-fold increase of the smear of Ty1 cDNA in 8-day-old stationary phase wild-type cells when compared to growing cells and ∼70-fold increase in *nuc1*Δ, which may explain the higher level of Ty1 insertions in *nuc1*Δ cells (**Fig. 4b**). Together, the increase of partially replicated retrotransposons in wild-type and particularly in *nuc1*Δ cells may contribute to inserted cDNA at DSBs. Enzymatic activities of purified EndoG nucleases show a preference for cleavage of ssDNA over dsDNA and the ability to degrade RNA in DNA:RNA hybrids ^34, 50, 51^, which could explain the strong impact of Nuc1 on intermediates of Ty replication, many of which are ssDNA or RNA:DNA hybrids. We propose that Nuc1 nuclease partially limits cDNA insertions by degrading incomplete Ty1 cDNA in stationary phase cells.

**Figure 4.**
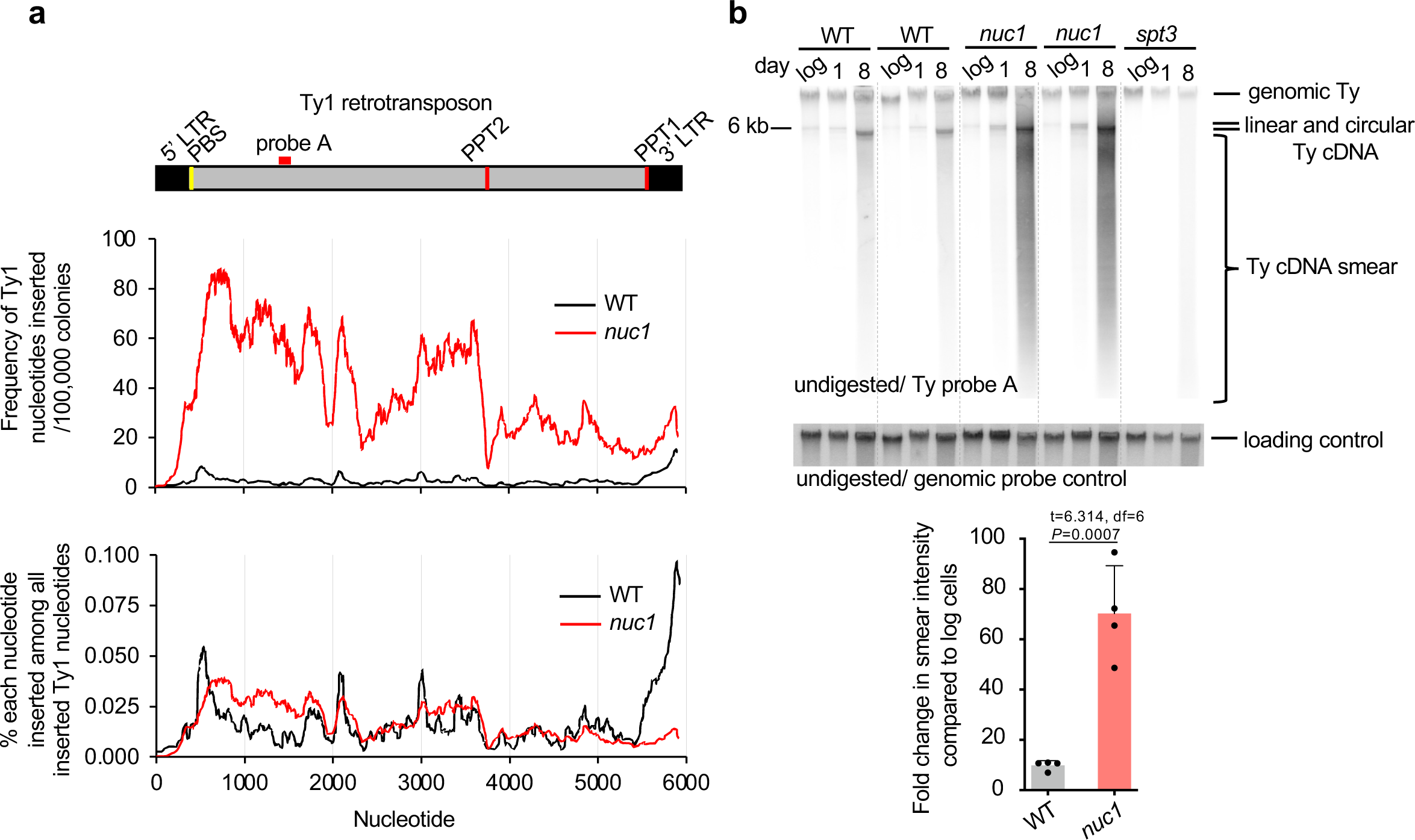
Nuc1 degrades retrotransposon cDNA replication intermediates. **a**, Analysis of Ty1 sequences inserted at DSBs in wild-type and *nuc1*Δ cells. Frequency of each Ty1 nucleotide insertion per 10^6^ of NHEJ products or among all Ty1 nucleotides inserted. Schematic of Ty1 shown at top indicating long terminal repeats (LTR), primer binding site (PBS) and polypurine tracts (PPT1/2). **b**, Southern blot analysis of Ty1 cDNA in wild-type, *nuc1*Δ and control *spt3*Δ cells. Quantification of Ty1 cDNA smear is shown (mean ± SD; n=4; P-values determined using unpaired two-tailed t-test). Position of the probe used is shown in **a**.

## Discussion

Cytoplasmic DNA must be degraded to limit sterile inflammation in metazoans. We provide evidence that degradation of long mtDNA and retrotransposons cDNA is important for nuclear genome instability. In wild-type cells grown in optimal conditions, the transfer of cDNA or mtDNA to nuclei is very rare. However, in aging or in stationary phase when cells experience stress and limited nutrients, transfer of these DNA species to the nucleus dramatically increases. Yeast EndoG is major regulator of this transfer. Elimination of Nuc1 increases partially replicated retrotransposon cDNA fragments and their insertions in the genome. In stressed cells transcription of transposons is upregulated ^22^, however, with limited nutrients it is likely that neither complete cDNA nor nucleocapsids are formed efficiently, leaving cDNA intermediates vulnerable for Nuc1 mediated degradation. Nuc1 absence also increases transfer of very long extrachromosomal mtDNA to the nucleus that form unstable circles. Up to ∼20 or 250-fold increase of the transfer of long *TRP1* marked mtDNA in *nuc1*Δ cells was observed during starvation in haploid cells or during sporulation in diploids, respectively. In yeast, any circular DNA devoid of centromeres shortens the lifespan of the cells ^52^; therefore, elimination of mtDNA released from mitochondria during sporulation or starvation produces healthier cells. MtDNA carries many repetitive sequences including microsatellite DNA, direct and inverted repeats, sequences forming G4 structures and sequences that may serve as efficient origins of replication in the nucleus. Therefore, nuclear mtDNA circles that are often present at a high copy number could instigate genome instability, recruit proteins needed for regular genome replication, and likely integrate into the genome as was shown for other circular DNA fragments in yeast and humans ^53, 54^. Nuclear circular mtDNA has not yet been studied in a systematic way in mammalian systems. MtDNA is excluded from sequencing analysis of circular DNA because of the inability to distinguish it from hundreds/thousands of regular mtDNA per cell. However, several pieces of evidence suggest that very long mtDNA can transfer to nuclei. Recent studies demonstrated many cases of multicopy mega-NUMTs inserted in the human genome ^55, 56^. Long NUMTs are also common in cancer cells^13^. Finally, nuclear extrachromosomal circular mtDNA was observed in nuclei of human pluripotent and embryonic stem cells ^57^. Opposite to transfer of long mtDNA, formation of short NUMTs decreases in *nuc1*Δ cells. However, the highest frequency of stable short NUMTs in the presence of Nuc1 is at least 1000-fold lower when compared to the spontaneous transfer of long mtDNA to the nucleus in the absence of Nuc1. Therefore, while the Nuc1 nuclease generates small mtDNA fragments that can be integrated into DSBs, it prevents the far more frequent nuclear transfer of very long mtDNA that also occasionally integrates into the genome (model in **Fig. 3c**). We propose that cytoplasmic nucleases such as EndoG or TREX1 in human and other organisms apart from their role in immune response also prevent genome instability caused by free DNA species.

## Acknowledgments

We thank Dr. Peter Thorsness, David Garfinkel, James Haber, Frank Madeo for the gift of plasmids or strains. We thank Dr. James Haber for critical reading of the manuscript, Dr. Mitch McVey for helpful suggestions, Alma Papusha for technical help. This work was funded by grants from the National Institute of Health (GM080600 and GM125650 to G.I, R01AG052507 and R01AG081347 to W.D. and R01GM138407 to K.C.).

## Author contributions

Y.Y. constructed strains, designed experiments including amplicon sequencing method, conducted most of the experiments, and analyzed data. X. W. and K. C. developed the software toolkits and bioinformatic pipelines to map and analyze insertions, and conducted most of statistical data integration; J.F. constructed strains, performed analysis of Ty1 transposition, Ty1 cDNA, mtDNA amount, and analyzed Ty1 cDNA insertions; G.I., Y.Y. and P.T. designed experiments to study *TRP1*-mtDNA transfer to nucleus; P.H. helped with data interpretation; W.D., R.Y., N.N., B.M. provided expertise on yeast aging and helped with isolation of old yeast cells and yeast lifespan analysis; G.I., Y.Y., X. W., and K.C. designed experiments, analyzed the data, and wrote the manuscript. J.F. edited the manuscript.

## Methods

### Yeast strains

All yeast strains used here to study insertions at HO endonuclease induced DSB are derivatives of JKM139 strain, a gift from James Haber, and are listed in **Supplementary Table 2**. All strains used to study the transfer of *TRP1*-mtDNA to the nucleus are derivatives of PTY44 strain, a gift from Peter Thorsness. In this system, a *TRP1* marker was inserted upstream of the *COX2* gene and *TRP1* marker carries no homology to the nuclear genome where *TRP1* is entirely deleted (*trp1*-Δ1). The *nuc1*-H138A mutants were constructed using the plasmid pCORE-UH (a gift from Francesca Storici). First, the *KlURA3*-hphMX double maker were integrated into *nuc1* locus. Then a fragment of *nuc1* with a point mutation was PCR amplified from the plasmid pESC-nuc1-H138A (a gift from Frank Madeo) and transformed into cells to replace *KlURA3*-hphMX. All mutations were confirmed by Sanger sequencing.

### Yeast growth, media and DSB induction

For amplicon sequencing analysis in normal conditions, cells were grown in YEPD (1% yeast extract, 2% peptone, 2% dextrose) o/n to saturation. Cells were washed twice in YEP-Raffinose (1% yeast extract, 2% peptone, 2% raffinose) media and inoculated in 10 ml of YEP-Raffinose and grown overnight. Once cells reached the density of 2×10^7^/ml, they were plated on YPE-Gal plates (1% yeast extract, 2% peptone, 2.5% agar, 2% galactose. Galactose was filter sterilized and added to the autoclaved media). Control plating on YEPD plates provided information on the number of viable cells. Old yeast mother cells isolated as described below were plated on YEPGal plates the same way. For all analyses in stationary phase cells, we followed a previously established protocol for yeast growth ^58^. Yeast cells were initially streaked out on YEPD plates and individual colonies were inoculated in 3 ml SC media (1.72g/L yeast nitrogen base without amino acids (USBiological, cat# Y2030), 2g/L amino acid mix (Bufferad, cat#S0051), 0.5% ammonium sulfate, 2% dextrose) and incubated overnight at 30 °C. The culture was diluted into fresh 50 mL SC media in 250 mL flasks to an OD_600_ of 0.1. Logarithmic growth phase control cells were plated when the culture reached 1×10^7^ cells/ml. Cells were incubated with shaking for 24 hrs (day 1), 3 days, 8 days, 16 days or as specified in individual experiments and plated on YEPGal plates for HO induction. Colonies were collected after 5 days of incubation for analysis of NHEJ products.

### Measurement of survival, respiration and NHEJ efficiency

To measure yeast cell survival (CFU), the cells were diluted and spread on YEPD plates. Colonies were counted 5 days after plating. To check the respiration capacity, colonies on YEPD plates were replica plated to YPEGlycerol plates, where fermentation is not possible. The number of colonies that failed to grow on YPEG out of all colonies replica plated from YEPD is the proportion of respiration deficient cells. To analyze efficiency of DSB repair by NHEJ, cells were plated on YEP-Gal (GAL10-HO induction) and YEPD (no HO induction). As only a small fraction of cells survive constant HO induction, typically 100,000 cells were plated on YEPGal plates and 100 cells on YEPD plates. Efficiency of NHEJ was calculated as the percentage of cells growing on YEPGal out of all cells plated on YEPD plates.

### Isolation of old yeast cells

Populations of old yeast mother cells were isolated as previously described ^59^ with the following modifications. 3×10^8^ yeast cells from a fresh log-phase culture in YEPD were treated with 3 mg of Sulfo-NHS-LC-Biotin (Thermo Scientific) and were used to seed a 1L culture in YEP-Raffinose. Seeded cultures were incubated with shaking at 30 °C for 15 hours to age biotin-labeled cells. The aged biotin-labeled cells were isolated using 1 mL Dynabeads Biotin Binder (Thermo Scientific). Three rounds of aging and sorting were conducted to obtain old mother cells with an average replicative age of around 20. Control young cells were obtained from last sorting and wash.

### Yeast replicative lifespan measurement

The yeast replicative lifespan was determined in microfluidic devices (AD-Chips, obtained from Innovative Biochips) with time-lapse imaging as described previously ^60, 61^. Briefly, filter-sterilized YEPD was loaded into the AD-Chip channels at 20 µL/min using 20-mL syringes (BD Biosciences) driven by a KDS-230 pump (KD Scientific); medium flow was subsequently set to 1 µL/min prior to cell loading. Yeast strains were grown to mid-log phase in filter-sterilized YEPD and diluted 1:20 in the same prior to manually loading cells into the AD-Chip. After cell loading, the medium flow was set to 5 µL/min for the duration of the experiment. The AD-Chip was placed in an environmental chamber set to 30 °C for the lifespan analysis. Multi-position timelapse imaging was performed with an EVOS FL Auto system (Thermo Fisher), using a 20× objective and transmitted light optics. Images (4 images encompassing 480 traps per microfluidic channel) were taken every 15 minutes for 65 hours. The timelapse image series were analyzed with ImageJ (National Institutes of Health) and cell divisions were manually counted for a minimum of 50 cells from each strain.

### Analysis of Ty1 transposition

Rates of Ty1 retrotransposition were estimated using strains carrying a Ty1-270*his3-AI* reporter^47^. Individual colonies of WT and *nuc1*Δ strains were inoculated in 5mL YEPD media overnight at 24°C. Overnight cultures were then diluted and cells were grown to 1×10^7^ cells/mL before plating ∼20-30 cells on YEPD plates. After 3-5 days of incubation at 24°C, ten individual colonies per strain were inoculated to 8mL SC media and incubated at 24°C overnight. 2×10^8^ cells of each culture were plated on 150×15mm plates of synthetic complete medium lacking histidine, and 100 cells were plated on YEPD plates to measure viability. The remaining Day 0 culture was used to inoculate fresh 8mL SC media to an OD of 0.1. The plating procedure was repeated on Day 3 and Day 8 without dilution to fresh SC. After growth at 24°C for ∼7-10 days, His^+^ colonies were counted. Statistical analysis of retrotransposition rates was performed with Drake estimator as previously described ^62^, normalizing the number of cells plated by multiplying 2×10^8^ cells by the number of cells that grew on YEPD plates/100. P-values were determined using the bootstrap resampling approach.

### Analysis and quantification of mtDNA, Ty1 linear, circular, and smear cDNA amounts

For analyses of Ty1 cDNA or mtDNA in stationary phase cells, yeast cultures were grown as described above. Cells from log phase, saturated, and 8 days stationary phase were collected by centrifugation, washed in sterile water and frozen. Genomic DNA was extracted as described previously ^63^ with modification. Cells were resuspended in 500uL extraction buffer (2% SDS, 100mM Tris-HCl pH 8.0, 50mM EDTA) and 5μL β-mercaptoethanol and 10μL 100T Zymolyase (5mg/mL) were added. Cells were lysed by 30-60 minutes incubation with shaking at 37 °C. DNA was extracted using a standard phenol/chloroform extraction and digested with RNaseA (0.1 mg/mL) overnight at 37 °C. For analysis of Ty cDNA, undigested DNA was run on a 1 % agarose gel and subjected to Southern blot analysis using a Ty1 specific DNA probe amplified by PCR using primers Typeak fw and Typeak rv. For analysis of mtDNA amount the same DNA was probed with a mtDNA specific DNA probe amplified by PCR using primers mtDNA F1 and mtDNA R2. See **Supplementary Table 3** for primer sequences.

Band intensities corresponding to probed DNA fragments were analyzed using ImageQuant software. All quantifications were done with a normalized amount of DNA in each lane. For undigested DNA, normalization was done with a genomic mixture of three probes amplified using primers ACT1-fw and ACT1 rv, TRA1 fw and TRA1 rv, TOM1 fw and TOM1 rv. All probes were gel purified using the Nucleospin Gel and PCR Clean-up kit (Macherey-Nagel, cat# 740609). The pixel intensities of the bands corresponding to linear, circular, or smear of Ty cDNA were subtracted by the corresponding background intensity observed in *spt3* mutant cells that do not generate cDNA. The pixel intensities of cross-hybridizing bands observed in the *spt3* mutant cells that do not generate cDNA were subtracted from the bands corresponding to linear, circular, or smear of Ty cDNA. For smear, all DNA below linear cDNA was measured. The amounts observed in saturated culture cells (day 0) and day 8 were compared to cDNA levels observed in wild-type growing cells. Three to four repeats were done. T-test was used to quantify P-values.

### Amplicon sequencing of repaired MAT locus by “*Break-Ins”*

Cells from an overnight saturated YEPD culture were washed twice with YEP-Raffinose, inoculated into 10 ml YEP-Raffinose and incubated overnight at 30 °C. When the density of the culture was ∼2×10^7^ cells/ml, ∼1×10^7^ cells were spread on a YEP-galactose plates and incubated at 30 °C for 5 days. Each culture was spread on 5-7 150×15 mm plates. Colony number was counted. All colonies were collected, pooled together and mixed vigorously. ∼80 μl cells were spun down and the genomic DNA was extracted using standard glass beads and phenol/chloroform. The genomic DNA was treated with RNase A overnight. The genomic DNA was dissolved with water and adjusted to a concentration of 10 ng/μl. The construction of the sequencing library was adapted from Illumina’s amplicon sequencing protocol #15044223 Rev. B. Two rounds of PCR were performed to construct the sequencing library. For each sample, 3.2 μl genomic DNA, 0.5 μl 10 μM primers and 12.5 μl KAPA HiFi HotStart ReadyMix (Roche, 7958927001) was used for the first round of PCR for a total volume of 25 μl. The forward primers are 66-mers that contain (5’ to 3’): 33 bases Illumina adapter sequence, 3 bases unique home index, and 30 bases targeting the *Mat***a** locus which is located 11 bp upstream of the HO cleavage site. The reverse primers are 66-mers that contain (5’ to 3’): 34 bases Illumina adapter sequence, 3 bases unique home index, and 29 bases targeting the *MAT***a** locus which is located 18 bp downstream the HO cleavage site. The following conditions were used for the first round of PCR: 95 °C for 5 min; 22 cycles of 98 °C for 20 s, 65 °C for 30 s, 72 °C for 3 min; 72 °C for 10 min. 18μl PCR product was purified with 18 μl AMPure XP beads (Beckman Coulter, A63880) and eluted with 52.5 μl 10 mM Tris pH 8.5. For the second round of PCR, 5 μl purified PCR product, 5 μl Nextera XT V2 Index (Illumina, FC-131-2001) primer N7xx, 5 μl index primer S5xx, and 25 μl KAPA HiFi HotStart ReadyMix were used. The total volume of PCR is 50 μl. The following conditions were used for PCR: 95 °C for 5 min; 8 cycles of 95 °C for 30 s, 55 °C for 30 s, 72 °C for 3 min; 72 °C for 10 min. 40 μl PCR product was purified with 48 μl AMPure XP beads and eluted with 27.5 μl 10 mM Tris pH 8.5. The DNA concentration of each sample was determined with Qubit dsDNA BR Assay Kit (ThermoFisher, Q32850) and the average size of the DNA was determined with TapeStation. The DNA was diluted to 4 nM using 10 mM Tris pH 8.5. An equal amount of DNA was pooled from ∼20 samples into the library. The pooled library and PhiX library (Illumina, FC-110-3001) were denatured with 0.2 M NaOH separately and diluted with pre-chilled HT1 to 12 pM. 540μl pooled library and 60 μl PhiX were mixed, incubated at 96 °C for 2 min and immediately placed in an ice-water bath for 5 min. The denatured combined library was loaded into the MiSeq Reagent Kit v3 (600 cycle) (Illumina, MS-102-3003). The cluster density was ∼1100 K/mm^2^.

### Analysis of *TRP1*-mtDNA transfer to the nucleus

To analyze the frequency of transfer of *TRP1*-mtDNA to the nucleus, cells were grown as described above in SC media and plated on Trp^-^ plates. Control plating on YEPD informed about the number of viable cells plated. Trp^+^ colonies were scored after 5 days incubation at 30 °C. Analysis was repeated 3 to 8 times. The frequency of nuclear *TRP1*-mtDNA was calculated as the number of Trp^+^ colonies divided by the number of viable cells plated. To analyze the stability of the nuclear *TRP1*-mtDNA, an initial Trp^+^ colony was streaked onto Trp^-^ plates and grown for 2-3 days at 30 °C. Resulting single Trp^+^ colonies were streaked on nonselective YEPD plates and grown for 2 days at 30 °C. Once colonies formed, the plates were replica plated on Trp^-^ plates and the number of Trp^+^ colonies were counted. To analyze the size of nuclear *TRP1*-mtDNA, we streaked the Trp^+^ colonies for singles on Trp^-^ plates and inoculated in 3 ml Trp^-^ minimal liquid media containing 25 μg/ml ethidium bromide and incubated overnight at 30° C on a roller drum as previously described ^64^. The growth in Trp^-^ minimal liquid media with ethidium bromide media was repeated once more, and the cells were streaked on Trp-plates. A single colony was used to inoculate 5 ml Trp^-^ minimal media, and cells were grown to saturation at 30° C. Cells were pelleted and DNA was isolated using standard glass beads and phenol/chloroform procedure. Isolated DNA was digested with *Pst*I restriction enzyme, separated on 0.8% agarose gel, and subjected to Southern blot analysis using PCR amplified 3’ *TRP1* fragment (∼300 bp on each side of the *Pst*I cleavage site). Probe labelling and hybridization was done as described above. Size was estimated based on marker DNA.

### Analysis of insertion of transformed DNA

To obtain the set of 84 bp, 100 bp, 150 bp, 200 bp, 300 bp 400 bp, 500 bp and 1 kb dsDNA, we PCR amplified the DNA using Q5 polymerase (NEB, M0491) and Lambda DNA (NEB, N3011S) as a template. All of the ends of this set of PCR products have AACA. DNA was purified using NucleoSpin PCR clean-up kit (Macherey-Nagel, cat# 740609), dissolved with ddH_2_O and adjusted to a concentration of 0.6 μM. To obtain the set of 24 bp, 34 bp, 44 bp, 54 bp, 64 bp, 74 bp, dsDNA, the reverse-phase cartridge-purified oligos ordered from Sigma-Aldrich were dissolved with annealing buffer (10 mM Tris, pH 7.5, 1 mM EDTA, 50 mM NaCl) to 200 μM. Equal amounts of complementary oligos were mixed, heated at 95°C for 5 min and cooled down at room temperature. Every 10 bp of this set of dsDNA has AACA. To analyze the insertions of transformed DNA, the cells were collected when the density reached ∼2 × 10^7^ cells/mL in YEP-Raffinose and washed twice with ddH_2_O. For each transformation, 2 × 10^8^ cells were mixed with 240 μl 50% PEG3350, 36 μl 1M lithium acetate, water, and 20 μl 100 μM 24 bp-74 bp dsDNA or 64 μl 0.6 μM 84 bp-1kb dsDNA to total 360 μl. The mixture was incubated at 30°C for 30 min followed by 42°C for 30 min. The cells were centrifuged and resuspended in ddH_2_O, spread on YEP-Galactose plates and incubated at 30°C for 5 days. To test insertion of transformed DNA, the amfiSure PCR Master Mix (GenDEPOT, P0311) was used for colony PCR with the following conditions: 94°C for 5 min; 35 cycles of 94°C for 30 s, 52°C for 30 s and 72°C for 2 min 30 s. To test the insertion of short DNA (<100bp), the primers mata-F3 and mat-Rw3 were used for PCR, and the PCR products were visualized on EtBr containing 3% agarose gel. To test the insertion of long DNA (>100bp), the primers mata-F and mat-Rw were used for PCR, and the PCR products were analyzed by electrophoresis (1.2% agarose in 1x TBE buffer) at 120V for 30min.

### Processing of raw reads from *Break-Ins* analysis

Paired-end reads were assigned to samples based on the Illumina indexes. Then the custom indexes were used to discharge reads for which the Illumina indexes and the matched custom indexes were not mapped to the same sample. Thereafter, the software BBDuk v38.46 at the BBMap website (https://sourceforge.net/projects/bbmap/) was used to remove PhiX reads with default parameters. Subsequently, low-quality reads were removed while high-quality reads were retained with the average sequencing quality score (Q score) of each read to be at least 25 for bases in the *MAT***a** region and 15 for bases in the rest of the region. To calculate the average Q score in each region of a read, the quality symbol for each base in the FASTQ file was converted to the Q-score based on the Quality Score Encoding table provided at the Illumina web site (https://support.illumina.com/help/BaseSpace_OLH_009008/Content/Source/Informatics/BS/Qu alityScoreEncoding_swBS.htm). We then calculated the mean value of the scores in each region of each read.

### Detection of reads carrying insertions of 10 bp or longer

We used the software PEAR v0.9.11^65^ to merge the forward and reverse reads with default parameters. The merged reads were aligned to the yeast genome (S288C, 2μ plasmid sequence was included). For each pair of reads that could not be merged, e.g., because of no overlap between sequences of the two reads, we aligned the two reads to the genome separately. We used the software BLASTN v2.8.1 ^66^ to perform the alignment. A pair of merged reads was defined as having an insertion if the merged read were aligned to the *MAT***a** sequence but with at least 10 bp insertion at the break point of the *MAT***a** sequence. For a pair of reads that could not be merged, we considered it as having an insertion if the insertion portions of the two reads were at the 3’ ends and aligned to different loci that are at least 10 bp and at most 3 kb away from each other on the same chromosome.

### Defining unique insertions

If there was no sequencing error, two pairs of sequencing reads derived from a unique insertion event would have the identical sequence, and thus be recognized as a sequencing duplication. To eliminate insertion duplicates that differ in sequence due to sequencing error, we performed additional analysis. We used a 60 bp region (11 bp in the *MAT***a** region followed by 19 bp in the insertion donor sequence region at both junctions) of each insertion to identify the sequencing duplications of a unique insertion. The reads that had identical or nearly identical junctions were considered as originating from the same insertion. Specifically, two pairs of reads that aligned to each other based on the 60 bp region with at least 95% identity, at least 95% coverage, and no more than 6 bp mismatches and indels were defined as duplications of one unique insertion.

We next further defined additional duplications based on the genomic locations at which the reads aligned. For each insertion detected at the induced break point at the *MAT***a** locus, the inserted sequence might come from a single genomic location (single-donor) or multiple genomic locations (multi-donor) in the genome. For two pairs of reads (2 reads for each sequencing pair and thus 4 reads for two pairs) for the same single donor, we define the distance between their mapped locations at the left side of the insertion site (d_1a_), the distance between their mapped locations at the right side of the insertion site (d_1b_), the distance between their mapped locations at the left side of the donor site (d_1c_), and the distance between their mapped locations at the right side of the donor site (d_1d_). The two pairs of reads are defined as a duplication of one unique insertion if d_1a_ + d_1b_ <= 6 bp and d_1c_ + d_1d_ <= 6 bp. For a n-donor insertion, we will define the four distances for each donor i as d_ia_, d_ib_, d_ic_, and d_id_ (i = 1, 2, …, n). The two pairs are defined as a duplication if (1) the sum of all d_ia_ and d_ib_ is no more than 10 and (2) the sum of all d_ic_ and d_id_ is no more than 10.

Furthermore, among the unique insertions defined in the above two steps, we retrieved each insertion that had less than 10 pairs of reads and had more than 300 bp of sequence in the *MAT***a** region plus the donor region. For any two of these retrieved insertions that had a sequence identity greater than 80% with each other, we further defined them as a duplication. Finally, we retrieved each unique insertion that had no more than 10 read pairs in one sample but had more than 500 read pairs in another sample sequenced in the same batch. The insertion in the sample that displayed no more than 10 read pairs was recognized as contamination and thus discarded.

At the end, each unique insertion might have multiple pairs of reads that represent sequencing duplications. We selected one pair of reads to represent each unique insertion. For this purpose, we gave high priority to the read pairs for which the forward and reverse reads had overlap and thus could be merged. Thereafter, between different read pairs that each could be merged, we provide high priority to the pairs that have higher sequencing quality. Finally, between different read pairs that have the same sequencing quality, we gave high priority to the pairs that each have large number of identical copies.

### Defining the locations of the donor sequences of insertions

For more than 90% of the insertions, the donor locations could be easily defined based on the mapping result of BLASTN. However, it is more complicated to define the donor locations for a few insertions that belong to one of two categories. For the first category, the insertion sequences were not mapped to any donor location by the BLASTN algorithm. We thus used the BLASTN-short algorithm, which allowed us to further map some of these insertions to donor locations. For the second category, each insertion could be mapped to multiple alternative donors due to the great sequence similarity between the alternative donors. In this scenario, each alternative donor sequence covered a large, but not the full, proportion of the insertion sequence. Among the alternative donors, we select the donor that covered the largest proportion of the insertion sequence. Finally, because each insertion sequence might consist of several different donor sequences, our algorithm counts the number of donors in each insertion.

### Analysis of trimming and microhomology around DSB junction

Trimming of *MAT***a** sequence near the junction of an insertion was common. Deletion was defined based on the missing nucleotides in the mapping result generated by BLASTN, which aligned the sequencing read to the original *MAT***a** sequence. The insertion was defined based on the extra nucleotides that could not be mapped to the *MAT***a** sequence and also could not be mapped to any sequence near the original donor location of the insertion. The microhomology between *MAT***a** and insertion donor was defined as sequence mapped to both the terminal region of the donor location and the terminal region of the *MAT***a** sequence flanking the break site. The microhomology sequence was defined by requiring perfect (100 % identity) match with both the donor sequence and the *MAT***a** sequence.

### Defining genomic features of the donor sequences of inserted DNA

We used Bedtools v2.253 to retrieve genomic features that overlap with or are located close to the locations of donor sequences in the genome. Locations of the confirmed origins of replication (ARSes) were collected from the OriDB database (http://cerevisiae.oridb.org/). We collected locations of reference R-loops from a published source ^67^. Known tandem repeats were downloaded from UCSC genome browser as a compacted file provided at the website (https://hgdownload.soe.ucsc.edu/goldenPath/sacCer3/bigZips/chromTrf.tar.gz). All other genomic features, i.e., tRNA, telomere, centromere, were acquired from the SGD database (https://downloads.yeastgenome.org/). To evaluate significance of association between a type of genomic feature and the donor locations of insertions, we performed comparison between the observed locations of the insertion sequences and a set of random control locations. For each genotype, random control locations were generated by shuffling the real observed locations of the insertions using the algorithm BEDTools shuffle ^68^. Therefore, the control locations were generated to have the same sizes as for the locations of the observed insertions.

Overlapped events between insertions and genome features are defined as having at least 1-bp overlap. Distance analysis was based on edge distance between an insertion and its closest genome feature. Insertions coming from Ty retrotransposons, mtDNA, rDNA, *MAT***a** or 2µ plasmid were excluded from these analyses. R-loops, ARS, telomeres, centromeres, tRNAs, and tandem repeats were used for comparison between the observed insertion locations and randomized control locations. Randomized control locations were shuffled by 100,000 times bootstrapping. P values for genomic feature overlap or proximity were calculated by one-sided permutation test.

### Frequency of Ty1, mtDNA, rDNA sequence insertions

All inserted sequences derived from the LTR retrotransposon (Ty1) were realigned further against the same Ty1 sequence (YPLWTy1-1). Due to the high divergence of Ty1 insertions, we requested a loose parameter for BLASTN realignments (-word_size 11 -evalue 0.01 -gapopen 5 -penalty -1 -perc_identity 60 -dust no -soft_masking false). Among these alternative alignments, we selected the one that covered the largest proportion of the insertion sequence. In the case that Ty1 inserts could map perfectly to both LTR regions, we selected the right LTR alignment.

Based on the aligned positions of Ty1 inserts, we measured the number of times each Ty1 nucleotide was inserted, *n*_t,i_. Frequency of nucleotide inserted at each Ty1 position were calculated per 100,000 independent NHEJ products as 100,000\**n*_t,i_ /*n* _colonies._ Percentage of each nucleotide inserted among all Ty1 insertions were measured by 100* *n* _t,i /_*n* _t, total._ *n*_t,i_ as the number of insertion events at the i position, whereas *n* _t, total_ is the total number of insertion times with all the Ty1 positions. And *n* _colonies_ is the total number of colonies. The same nucleotide frequency methods but without realignments were applied for inserts from mtDNA and rDNA.

### Data and code availability

The *Break-ins* data has been archived at GEO under the accession number [GSE246469]. All original code has been deposited on GitHub. The code for insertion detection can be accessed at the following GitHub repository: https://github.com/gucascau/iDSBins.git. For short indel detection, the code can be accessed at this GitHub repository: https://github.com/gucascau/iDSBindel.git. The features related to insertion analysis have been uploaded to a dedicated GitHub repository, which can be found here: https://github.com/gucascau/LargeInsertionFeature.git. Any additional information required to reanalyze the data reported in this paper is available from the lead contact upon request.

## Supplementary Figure Legends

**Supplementary Figure 1.**
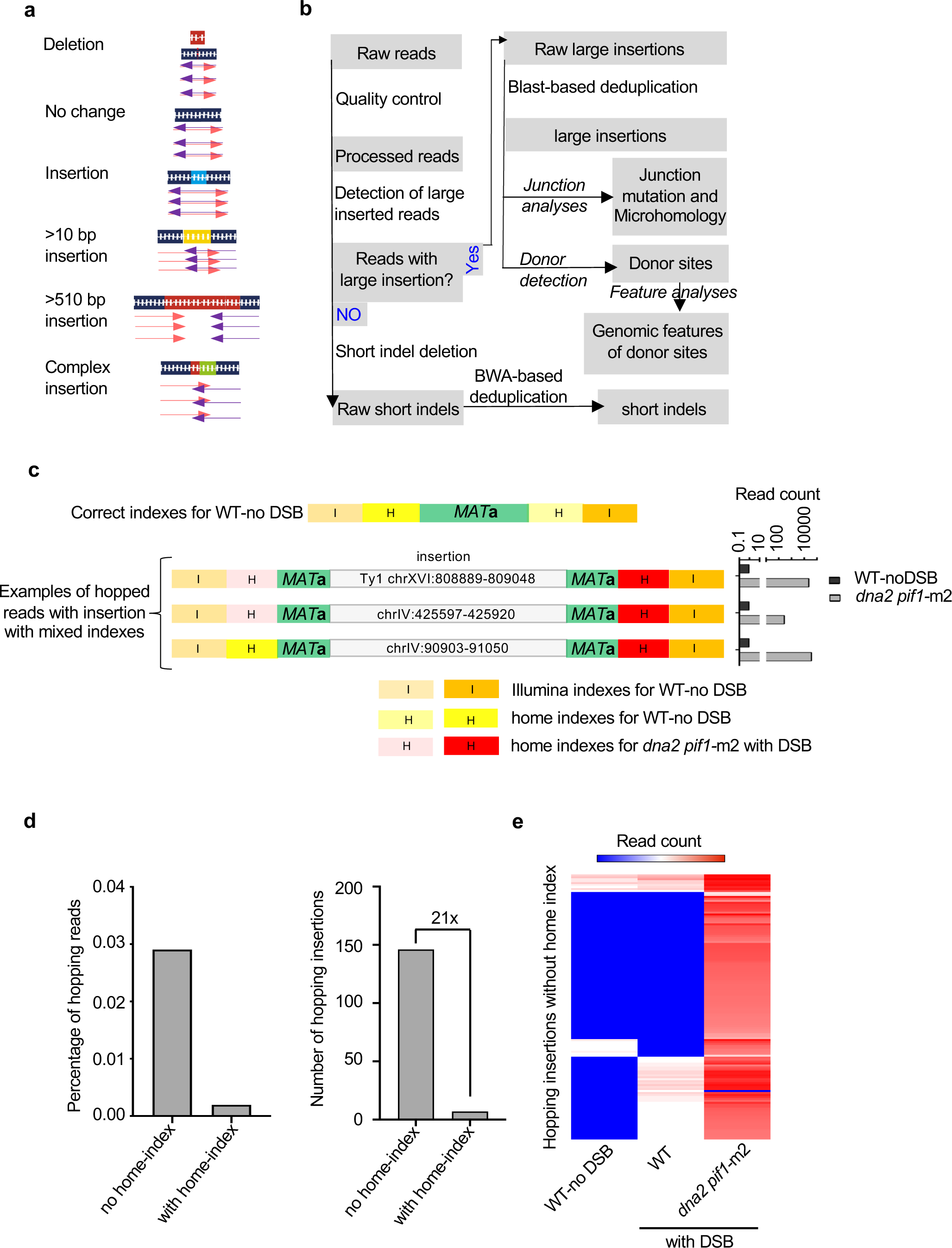
*Break-Ins* method for analysis of templated insertions. **a,** Schematic showing different types of sequence variation at DSB. **b**, Flowchart showing the computational pipeline for detecting different types of sequence variation at DSB. **c**, An example of cross-sample contamination indicated by index hopping during amplicon grigosequencing by MiSeq. Three examples show reads that have Illumina indexes specific for the DNA sample from a wild-type no DSB control but carry insertions. These reads can be eliminated by analysis of secondary home indexes. Number of false insertion reads in wild type compared to same insertion reads in *dna2*Δ *pif1-*m2 is shown on the right. **d**, Comparison of read hopping identified based on MiSeq amplicon sequencing with and without home indexes. **e**, Number of hopped reads without home index in single sequencing run.

**Supplementary Figure 2.**
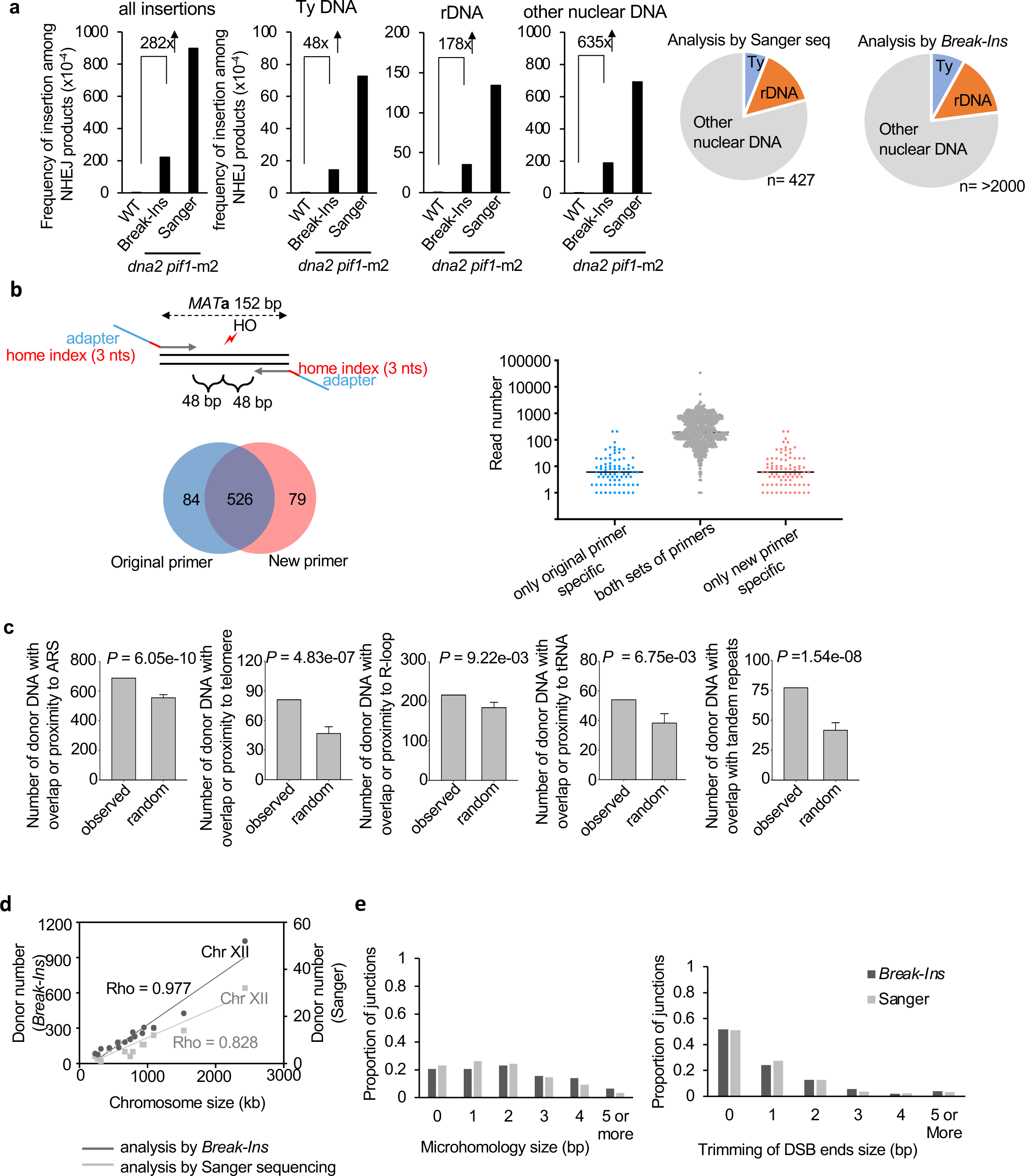
Comparison of Sanger sequencing and *Break-Ins* analysis. **a**, Frequency and types of DNA inserted at DSBs in wild-type or *dna2*Δ *pif1*-m2, (n – number of NHEJ products tested in shown in Supplementary Table 1). Sanger sequencing data are taken from previous publication^21^. **b**, Comparison of *Break-Ins* analysis done with two different primer sets. Scheme showing primer position with respect to DSB ends and number of insertions identified by both or just one set of primers (left panel). Original primer set is shown in Figure 1b. Most of the unique insertions are represented by low read number (right panel). **c**, Analysis of features of DNA inserted from nuclear genome at DSBs in *dna2*Δ *pif1-*m2. P-values were calculated using one-sided permutation test. Proximity is defined as sequence within 1 kb from ARS or telomere and within 0.2 kb from tRNA or R-loop. **d**, Distribution of donor DNA per chromosome. **e**, Analysis of microhomology and DSB ends trimming at insertion junctions in *dna2*Δ *pif1-*m2.

**Supplementary Figure 3.**
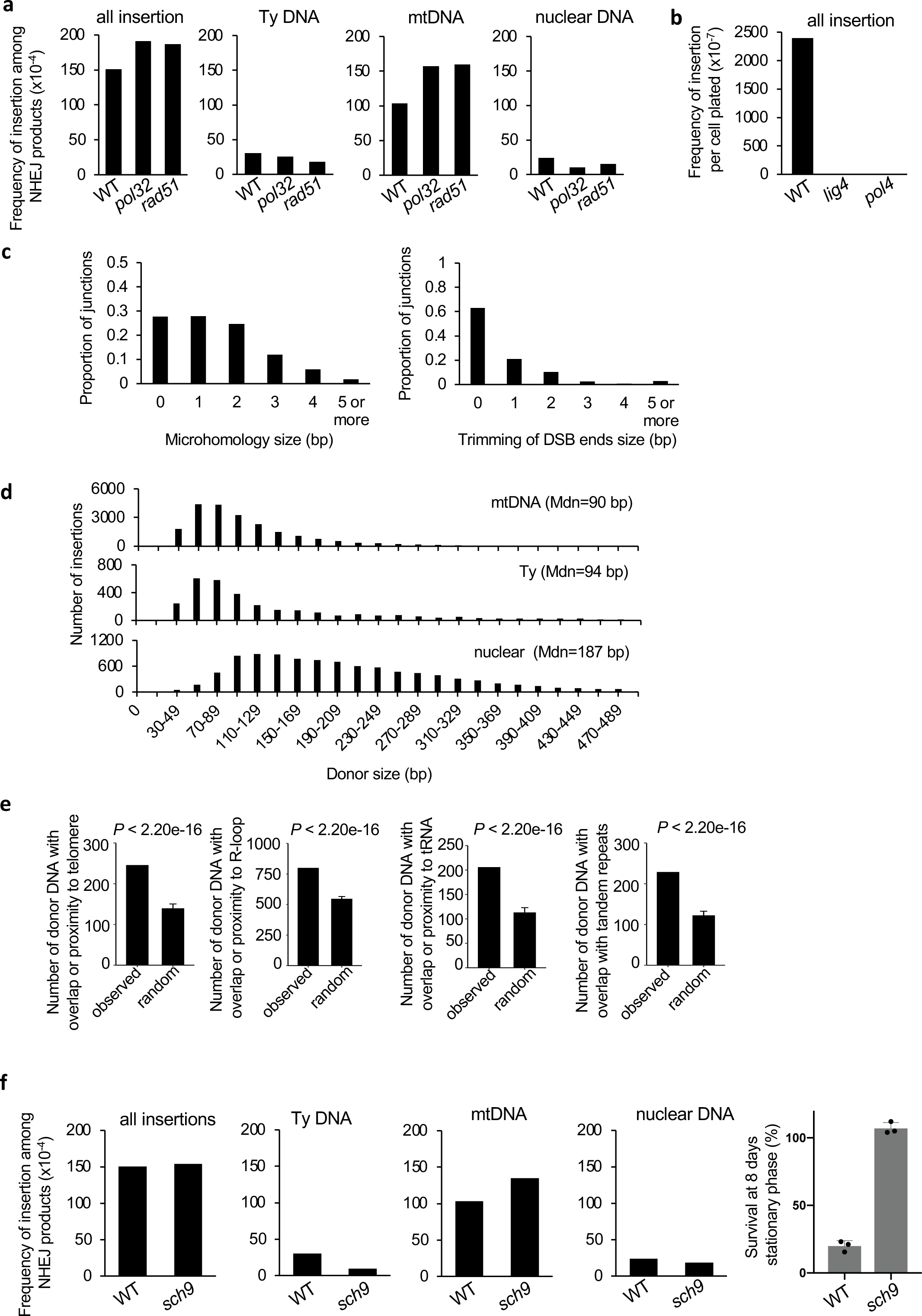
Analysis of templated insertions in mutants affecting DNA repair and aging. **a**, Frequency and types of DNA inserted at DSBs in *rad51*Δ and *pol32*Δ, (n - number of NHEJ products analyzed in shown in Table S1). **b**, Frequency and types of DNA inserted at DSBs in *lig4*Δ and *pol4*Δ. **c**, Analysis of microhomology and DSB ends trimming at insertion junctions in 16 days wild-type stationary phase cells. **d**, Insertion size analysis originating from mtDNA, Ty DNA, and nuclear genome. **e**, Analysis of features of DNA inserted from nuclear genome at DSBs in wild-type stationary phase cells. Insertions observed in all days (3, 8, 16) were combined for this analysis. P-values were calculated using one-sided permutation test. Proximity is defined as sequence within 1 kb from ARS or telomere and within 0.2 kb from tRNA or R-loop. **f**, Frequency and types of DNA inserted at DSBs in *sch9*Δ at 8 days in stationary phase (n-number of NHEJ products analyzed in shown in Supplementary Table 1). Viability of wild-type and *sch9*Δ cells at 8 days in stationary phase is shown on the right.

**Supplementary Figure 4.**
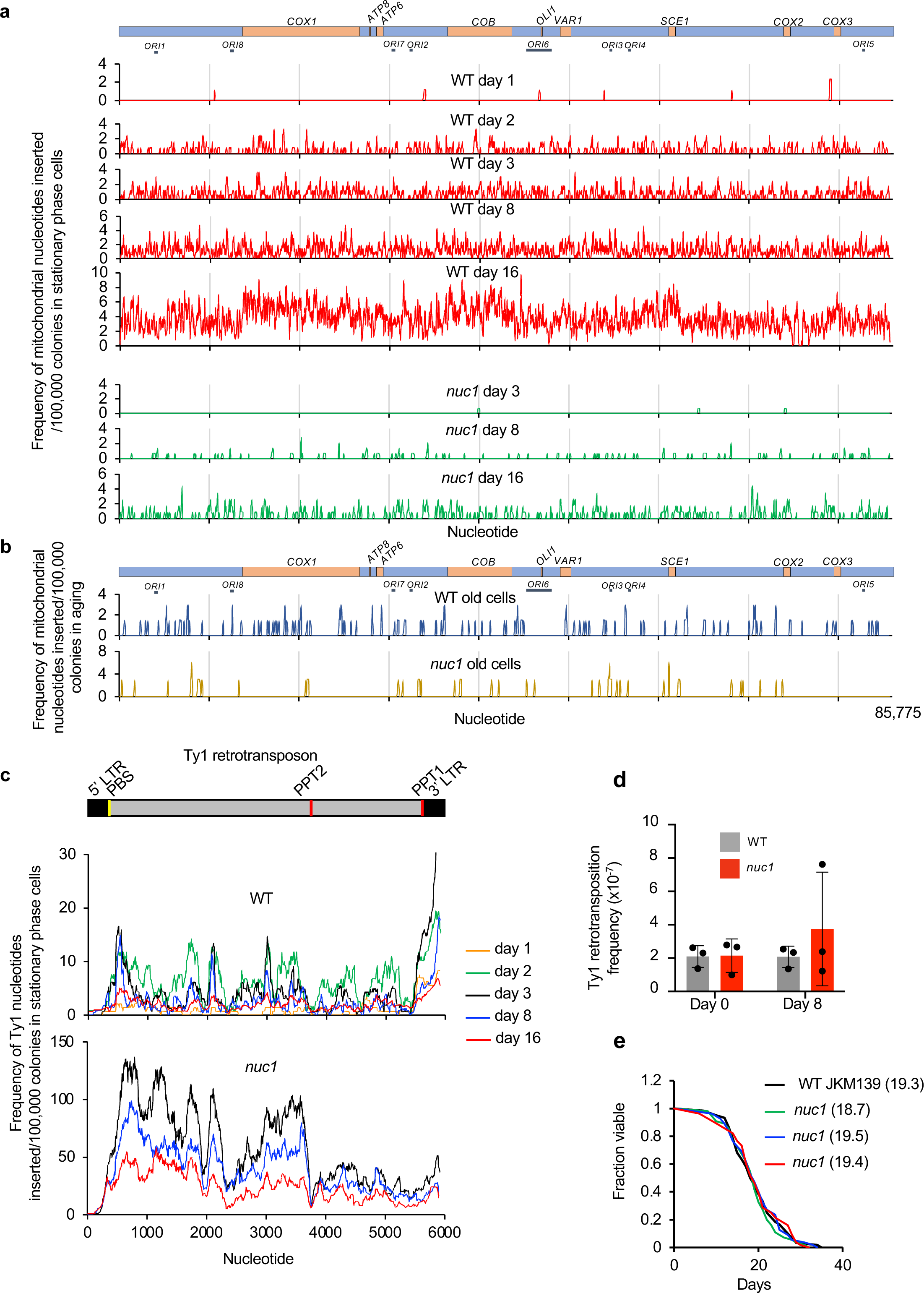
Analysis of mtDNA and Ty1 sequences inserted at DSB. **a**, Analysis of sequences inserted at DSB from mtDNA in wild-type and *nuc1*Δ during stationary phase. **b**, Analysis of sequences inserted at DSB from mtDNA in wild-type and *nuc1*Δ in aged cells. **c**, Analysis of sequences inserted at DSB from Ty1 in wild-type and *nuc1*Δ during stationary phase. **d**, Transposition frequency in wild-type and *nuc1*Δ mutant in stationary phase cells, (mean ± SD; n≥3; P values determined using unpaired two-tailed t-test). **e**, Analysis of lifespan of wild-type and three independent *nuc1*Δ mutant cells. Average lifespan in each strain is shown in parentheses.

**Supplementary Figure 5.**
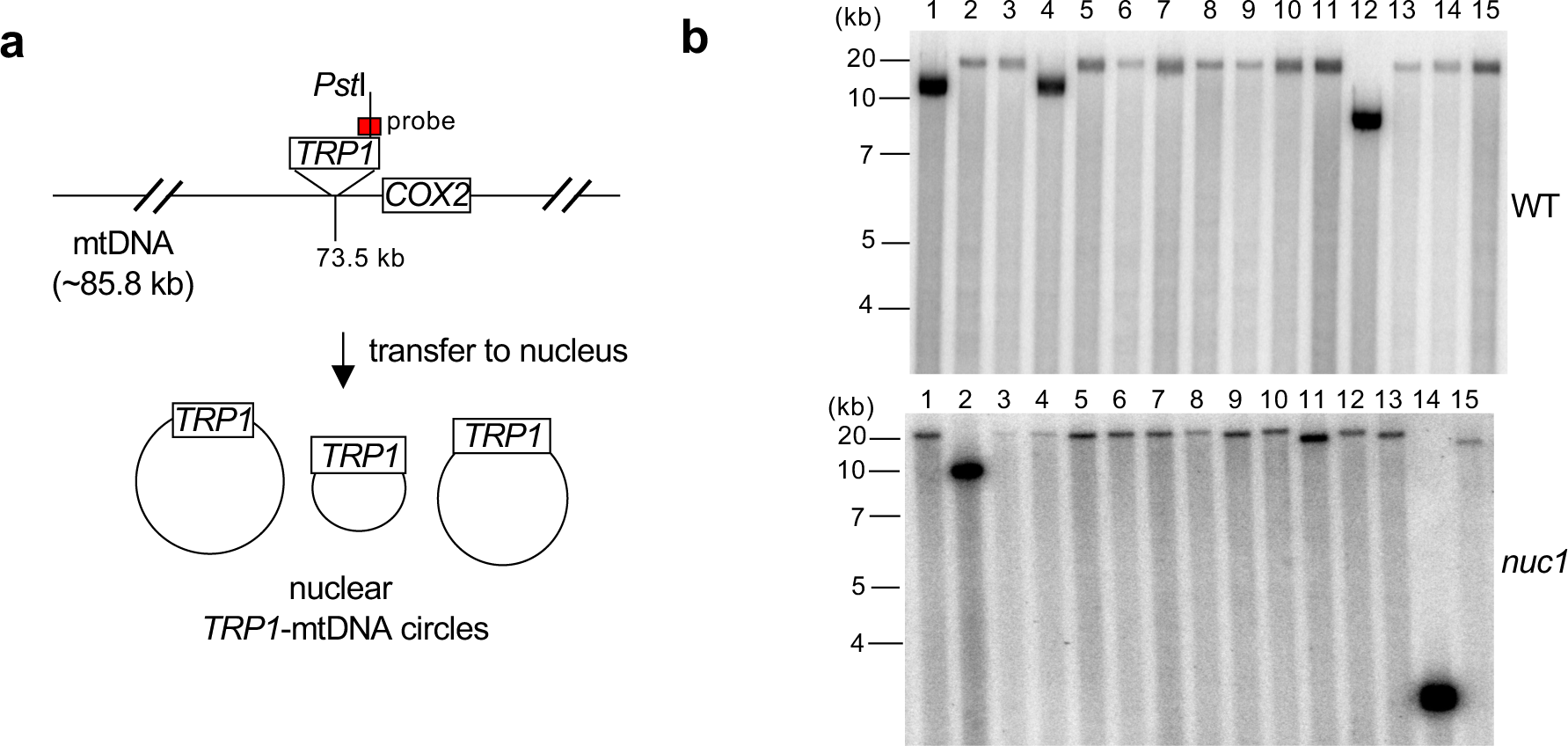
Analysis of nuclear *TRP1*-mtDNA. **a**, Schematic of mtDNA marked with *TRP1* and circular *TRP1*-mtDNA transferred to nucleus. DNA was digested with the *Pst*I restriction enzyme that cuts once 3’ to the *TRP1* reporter inserted within mtDNA but not anywhere within the mtDNA itself. Southern blot analysis with a *TRP1*-specific probe is expected to show two fragments if mtDNA*-TRP1* were linear and one fragment if it were circular. **b**, Southern blot analysis of nuclear *TRP1*-mtDNA digested with *Pst*I in wild-type and *nuc1*Δ cells. DNA probe location is indicated in **a**. 15 independent *TRP1*-mtDNA for each were analyzed. A single band was observed in all cases.

